# High-content single-cell combinatorial indexing

**DOI:** 10.1101/2021.01.11.425995

**Authors:** Ryan M. Mulqueen, Dmitry Pokholok, Brendan L. O’Connell, Casey A. Thornton, Fan Zhang, Brian J. O’Roak, Jason Link, Galip Gurkan Yardmici, Rosalie C. Sears, Frank J. Steemers, Andrew C. Adey

**Affiliations:** Oregon Health & Science University, Department of Molecular and Medical Genetics, Portland, OR; ScaleBio, CA; Oregon Health & Science University, Cancer Early Detection Advanced Research Center, Portland, OR; Oregon Health & Science University, Knight Cancer Institute, Portland, OR; Oregon Health & Science University, Department of Oncological Sciences, Portland, OR; Oregon Health & Science University, Brendan Colson Center for Pancreatic Care, Portland, OR; Oregon Health & Science University, Knight Cardiovascular Institute, Portland, OR

## Abstract

Single-cell genomics assays have emerged as a dominant platform for interrogating complex biological systems. Methods to capture various properties at the single-cell level typically suffer a tradeoff between cell count and information content, which is defined by the number of unique and usable reads acquired per cell. We and others have described workflows that utilize single-cell combinatorial indexing (sci)^1^, leveraging transposase-based library construction^2^ to assess a variety of genomic properties in high throughput; however, these techniques often produce sparse coverage for the property of interest. Here, we describe a novel adaptor-switching strategy, ‘s3’, capable of producing one-to-two order-of-magnitude improvements in usable reads obtained per cell for chromatin accessibility (s3-ATAC), whole genome sequencing (s3-WGS), and whole genome plus chromatin conformation (s3-GCC), while retaining the same high-throughput capabilities of predecessor ‘sci’ technologies. We apply s3 to produce high-coverage single-cell ATAC-seq profiles of mouse brain and human cortex tissue; and whole genome and chromatin contact maps for two low-passage patient-derived cell lines from a primary pancreatic tumor.

## Main

The core component of many sci-assays, as well as ATAC-seq, is the use of transposase-based library construction. While the transposition reaction itself (tagmentation) is highly efficient, viable sequencing library molecules are only produced when two different adaptors, in the form of forward or reverse primer sequences, are incorporated at each end of the molecule. This occurs only 50% of the time (Figure 1a, left). To combat this inefficiency, strategies including the use of larger complements of adaptor species^3^, incorporation of T7 promoters to enable amplification via *in vitro* transcription^4–6^, or reverse adaptor introduction through targeted^7^ or random priming^8^ have been developed; however, these methods are often complex and result in limited efficiency improvements. Here, we present a novel means of adapter replacement to produce library molecules tagged with both forward and reverse adaptors for top and bottom strands, overcoming this efficiency bottleneck. This format permits the use of a DNA index sequence embedded within the transposase adaptor complex, enabling single-cell combinatorial indexing (sci) applications, where two rounds of indexing are performed — the first at the transposition stage, and second at the PCR stage^1,9,10^.

**Figure 1.**
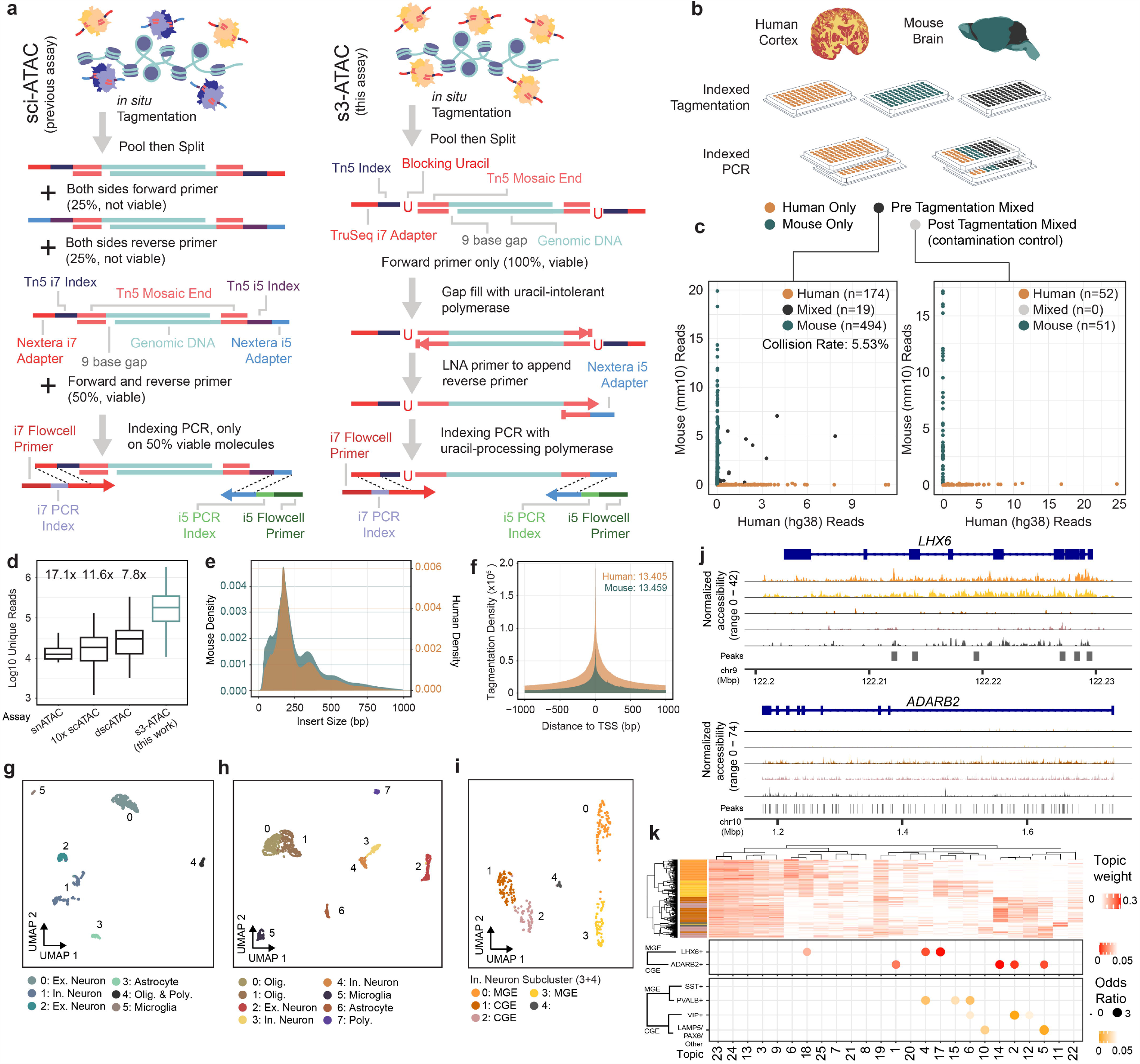
Symmetrical strand single-cell combinatorial indexing ATAC-seq (s3-ATAC). (a) Schematic of standard sci-ATAC library construction (left). Schematic of s3-ATAC library construction with intermediate steps of adapter switching leading to increased genomic molecule capture rate (right). (b) Experimental flow through and plate layout for the mixed-species experiment, including tagmentation and PCR plate conditions per well.(c) Point plots of single cell libraries with counts of unique reads aligned to mouse or human chromosomes in a chimeric reference genome. Points are colored to reflect species assignment (see Methods) in both pre-tagmentation mixing (left) and post-tagmentation mixing (right). (d) Comparison of library complexity for s3-ATAC mouse whole-brain sampled cells to previously reported data sets. All comparisons to our data are significantly less (Welch’s two-sample t-test, p value <0.01). Fold improvement of our library complexity per method is listed above the method^14–16^. (e) Insert size distribution of human and mouse libraries reflexing nucleosome banding.(f) Enrichment of reads at transcription start sites (“TSS”) for human and mouse libraries with enrichment calculation following ENCODE standard practices. (g) UMAP projection of mouse whole brain cell samples (n=837 cells) colored by cluster and cell type assignment. (h) UMAP projection human cortex cell samples (n=2,175 cells). (i) Subclustering and UMAP projection of human cortical inhibitory neurons (clusters 3 and 4 from panel h., n=342) (j) Genome coverage track of human inhibitory neurons (n=342) aggregated over 5 subclusters for genomic locations overlapping MGE and CGE marker genes *LHX6* and *ADARB2*, respectively.(k) Hierarchical clustering of topic weight per cell (top). Hypergeometric test of gene set analysis enrichment for human inhibitory neuron marker genes (bottom; Fisher’s exact test, see Methods).

Our strategy, <u>s</u>ymmetrical <u>s</u>trand <u>s</u>ci (s3; Figure 1a, right) uses single-adapter transposition to incorporate the forward primer sequence, the Tn5 mosaic end sequence and a reaction-specific DNA barcode. As with standard tagmentation workflows, extension through the bottom strand is then performed to provide adaptor sequences on both ends of each molecule; however, the s3 transposome complexes contain a uracil base immediately following the mosaic end sequence. Use of a uracil-intolerant polymerase therefore prevents extension beyond the mosaic end into the DNA barcode and forward adaptor sequence. A second template oligo is then introduced that contains a 3’-blocked locked nucleic acid (LNA) mosaic end reverse complement sequence with a reverse adaptor sequence 5’ overhang. This oligo favorably anneals to the copied mosaic end sequence, due to the higher melting temperature of LNA, and acts as a template for the library molecule to extend through and copy the reverse adaptor. This results in all library fragments having both a forward and reverse adaptor sequence. The LNA-templated extension is carried out over multiple rounds of thermocycling to ensure maximum efficiency of reverse adaptor incorporation. Furthermore, adapter sequences are designed such that standard sequencing recipes can be used instead of the custom workflows and primers that are required for current indexed transposition technologies^11,12^.

We first sought to establish the s3 technique to assess chromatin accessibility. In s3-ATAC, nuclei are isolated and tagmented using our single-ended, indexed transposomes and carried through the adaptor-switching s3 workflow (Figure 1a). To ensure we attain true single-cell libraries without contamination from other nuclei, and minimal barcode collisions, we performed a mixed-species experiment on primary frozen human cortical tissue from the middle frontal gyrus and frozen mouse whole brain tissue (Figure 1b). We elected to perform this test on primary tissue samples instead of an idealized cell line setting to more accurately capture the rates of cross-cell contamination. Levels of crosstalk were assessed at both points of possible introduction: the tagmentation and PCR stages; by mixing nuclei from the two samples before tagmentation as well as after. Additionally, pure species libraries were produced by leveraging the inherent sample multiplexing capabilities of sci workflows. In the experimental condition where nuclei were mixed prior to any processing, *i*.*e*. pre-tagmentation, we observed a total estimated collision rate of 5.53% (Figure 2dc; 2 × 2.77% detected human-mouse collisions), comparable to existing methods and tunable based on the number of nuclei deposited into each PCR indexing reaction. Zero collisions were observed in the post-tagmentation experimental conditions, suggesting no molecular crosstalk during s3 adapter switching or PCR.

**Figure 2.**
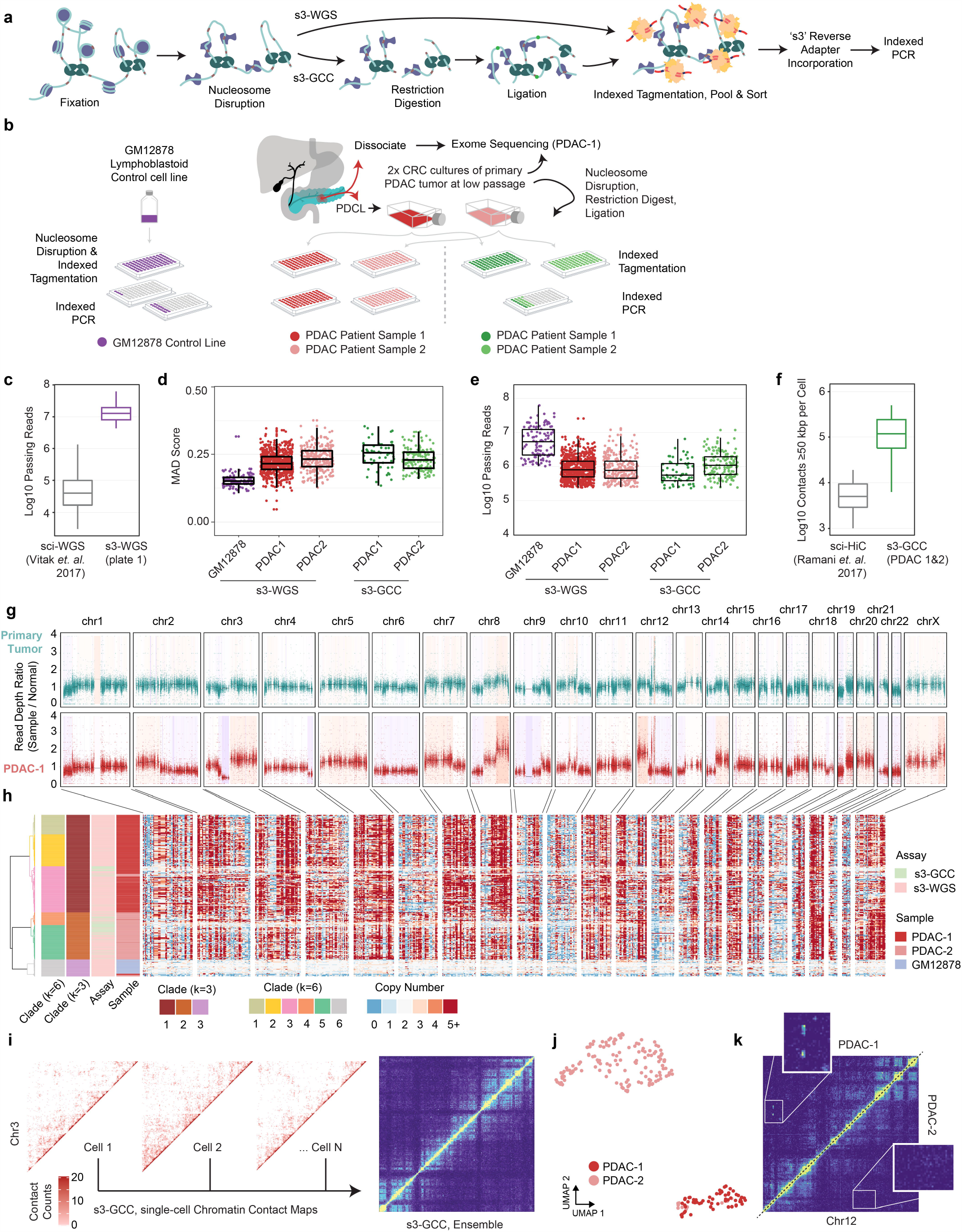
s3 whole genome sequencing (s3-WGS) and genome conformation capture (s3-GCC). (a) Schematic of sci-WGS and sci-GCC library construction. (b) Experimental flow through and plate layout for PDAC and control diploid line. (c) Boxplot of read count per cell for matched GM12878 cell line^9^. (d) Boxplot of MAD score per cell per sample and assay. (e) Boxplot of reads passing filter per cell. (f) Comparison boxplot of s3-GCC and sci-HiC distal contacts (≥50kbp) per cell^33^. (g) Whole exome sequencing of the primary tumor and PDCL. Scatterplot of reads per bin with a shading of called copy number variation. (h) Single-cell whole genome copy number calling on 500 kbp bins genome-wide. Cells (rows) are hierarchically clustered and annotated by assay, sample, and assigned clade (left). (i) Representative single-cell contact maps (raw counts) at 1 Mbp resolution for chromosome 3 and ensemble contact map profile at 500 kbp resolution. (j) scHiCRep dimensionality reduction and clustering of single-cell distal contact profiles. (k) Subclonal translocation on chr12 specific to PDAC-1.

In total, we generated 2,175 human and 837 mouse single-cell ATAC-seq profiles passing quality filters (Methods) across four PCR indexing plates (Figure 1b). We then assessed the total unique sequence reads obtained per cell as a function of the total aligned reads, *i*.*e*. the library complexity. One of our mixed species plates was sequenced to beyond 50% saturation (duplicate reads / total reads), to represent the sequencing depth obtained where diminishing returns of increased sequence depth become excessive^9^. For the mouse cells, the mean sequencing saturation per cell was 63.6% and resulted in a median unique read count per cell of 178,069 (mean = 258,859). The human cells reached a mean sequencing saturation of 56.6% with a median unique reads per cell of 99,882 (mean = 175,361). We additionally sequenced a plate that contained only human cells to a mean sequencing saturation of 70.4% which produced a median of 100,280 (mean = 146,937) unique reads per cell. When compared to other single-cell ATAC-seq datasets performed on mouse whole brain tissue, our mouse s3-ATAC libraries contain substantially greater reads per cell with 17.1x, 11.6x and 7.8x fold improvement compared to snATAC, 10X Genomics scATAC, and dscATAC, respectively (Figure 1d)^14–16^. Read count increases can be indicative of poor ATAC-seq library quality, with increased depth reflecting increased noise and loss of signal at open chromatin regions. To address this, we first assessed read pair insert sizes, revealing the characteristic nucleosome-size banding distribution of ATAC-seq (Figure 1e)^17^. We next calculated transcription start site (TSS) enrichment using the approach defined by the ENCODE project (Methods). This produced significant enrichment for both species at 13.4 for human, well above the ‘ideal’ standard (>7) and 13.5 for mouse, within the acceptable range and just below ideal (>15). Similarly, the fraction of reads in pile-up genomic regions (“peaks”; FRiP) was comparable to other single-cell ATAC technologies at 31.95% and 29.15% as measured using 292,156 and 174,653 peaks for human and mouse cells respectively. However, FRiP is largely dependent on the number of peaks called, which influenced heavily by cell number and total sequence depth obtained. When expanding to a human cortex high-depth ATAC-seq peak set, a mean of 48.1% of reads were present in peaks, and mean of 78.2% of reads for mouse cells using a high-depth mouse brain ATAC-seq peak set (Methods).

With ample signal, we next sought to discern cell types present within the complex tissues. For each species, we used peaks called on aggregate data to construct a count matrix followed by dimensionality reduction using the topic-modeling tool *cisTopic*^18^ which we then visualized using UMAP^19^, performed graph-based clustering at the topic level, and processed via *Signac*^20^. Clear separation of cell types was observed using marker gene signal and differential accessibility profiles (Figure 1g-h, Supplementary Figure 1)^16,21^. Notably, even with the modest cell count produced by this experiment, the quality improvements allow us to interrogate subclusters of inhibitory neurons previously difficult to distinguish in atlas-level datasets (Figure 1i)^22^. With our improved cell depth, we were able to discern caudal and medial ganglionic eminence inhibitory neurons by marker gene coverage plots across 342 GAD1+ cells (“CGE” and “MGE”, respectively; Figure 1j). From these, we identified 157 GAD1+, ADARB2+ CGE cells and 168 GAD1+,LHX2+ MGE cells. We separated 17 cells (subcluster 4) with apoptotic stress markers likely due in part to post-mortem sampling, which could potentially compound common single cell ATAC analyses. Aggregated genomic signal over our Topic-based dimensionality reduction was used to support our marker gene cell subtype discrimination and describe differentially accessible loci in human cortical inhibitory neurons (Figure 1k).

We then extend the improvements in data quality produced by s3-ATAC to other sci-workflows. This includes our previously-described sci-DNA-seq method^9^ that produces single-cell whole genome sequencing libraries (s3-WGS) and a novel strategy to incorporate the core components of HiC library preparation but without ligation junction enrichment to produce whole <u>g</u>enome and <u>c</u>hromatin <u>c</u>onformation information (s3-GCC; Figure 2a). Both strategies disrupt nucleosomes to acquire sequence reads uniformly across the genome^9^. We first tested s3-WGS by producing two small-scale libraries on the euploid lymphoblastoid cell line, GM12878. The first library comprised only four wells at the PCR stage for a target of 60 cells, allowing us to sequence the library to high depth (Figure 2b). This produced a median passing read count per cell of 12,789,812 (mean = 15,238,184), across 45 QC-passing cells (75% cell capture efficiency). With our sequenced library at 72.35% saturation; our complexity is notably higher than the predecessor sci-DNA-seq technology which produced a median of 43,367 reads per cell (mean = 103,138) at the same sequencing saturation (295 and 148 fold improvement in median and mean, respectively; Figure 2d)^9^. The second preparation performed comparably, though sequenced to a lower total depth (15.98% saturation). We also confirmed that the coverage was uniform by assessing the median absolute deviation (MAD) across 500 kbp bins, which fell within 0.152 ± 0.025 (mean ± s.d.), comparable to other single-cell genome sequencing techniques (Figure 2d)^9,23,24^.

We performed s3-WGS and s3-GCC on two cultures of a cell line derived from a primary pancreatic ductal adenocarcinoma (PDAC) tumor (Figure 2b). PDAC is a highly-aggressive cancer that typically presents at an advanced stage, making early detection and study of tumor progression key^25^. PDAC studies suffer from a low cancer cell fraction, thus we used a patient derived cell line (PDCL) generated directly from tumor and maintained at fewer than 10 passages. This method allows for multiple modalities of characterization and perturbation, while maintaining the heterogeneity present in the tumor sample^26^. We profiled two PDCL cultures (referred to as PDAC-1 and PDAC-2) to capture variance that may arise during passaging from a parent line derived from a tumor harboring a driver mutation in the oncogene *KRAS* (p.G12D). For our s3-WGS preparations, we produced 773 and 256 single-cell libraries with a mean passing read counts of 1,181,128 and 1,299,949 for PDAC-1 and 2 (at a combined median of 28.46% saturation), respectively. The s3-GCC libraries contained 57 and 145 cells produced a mean passing read count of 973,397 and 1,588,926 (combined median 73.25% sequencing saturation) for PDAC-1 and 2, respectively (Figure 2e). MAD scores for the two lines were greater than that of the euploid karyotype of GM12878, 0.219 ± 0.041 (mean ± s.d.); however, this is expected given the widespread copy number alterations present in the samples. In addition to the WGS component, the s3-GCC libraries also contained reads that were identified as chimeric ligation junctions that provide HiC-like chromatin conformation signal. Across both samples, we identified a mean of 118,048 reads per cell that capture genomic contacts at least 50 kbp apart from one another, a 14.8-fold improvement over the previous single-cell combinatorial indexing technique, sci-HiC^13^ (Figure 2f). Read pairs spanning ≥50 kbp accounted for a median of 15.6% and 17.0% of the total reads obtained per cell, which equates to an enrichment of 361- and 402-fold over that of the s3-WGS libraries for PDAC-1 and 2, respectively.

We first focused our analysis on the s3-WGS and the WGS component of the s3-GCC libraries to examine the copy number alterations present. To get a sense of the genomic landscape, we first performed copy number calling on whole exome sequencing (WES) libraries that were generated using primary tumor tissue and in bulk on the PDCL line (Figure 2g). This revealed a profile of copy number aberrations at finer resolution, with a more pronounced profile in the PDCL sample, likely due in part to the absence of euploid stromal cell contamination. We then processed all single-cell libraries using *SCOPE*^23^ which revealed a highly altered genomic landscape within each of the two samples. In line with paired karyotyping and bulk exome data, we see a similar pattern per cell of multi-megabasepair copy number aberrations when performing breakpoint analysis on 500 kbp windows, with a median depth per window of 81 reads. Using the inferred copy number profile within genomic windows for the three samples, GM12878 and the two PDCL cultures, we performed hierarchical and K-means clustering on the Jaccard distance between cell breakpoint copy numbers at two different centroid counts. For our optimal centroid value, we found a relatively clean separation between cell lines (k=3), for subclonal analysis we used a higher centroid count at a local optima (k=6). s3-WGS and s3-GCC cells cluster dependent on PDCL culture, reflecting our ability to capture genome-wide copy number data in our s3-GCC libraries (Figure 2h). We generated pseudo-bulk clades from the single-cell read count bins, with an average of 211.3 cells per clade and an average read count of 3,750 per 50 kbp bin. This revealed multiple fixed and subclonal genomic arrangements (Supplementary Figure 2). In PDAC-1 and PDAC-2 we see shared copy number loss of tumor suppressor genes *CDKN2A, SMAD4* and *BRCA2*^25,27^. In PDAC-2 we observed a subclonal amplification of *PRSS1*, a mutation that was fixed within our sampling for PDAC-1 and is associated with tumor size and a higher tumor node metastasis (TNM) stage^28^. This suggests that while the lines have the same origin, each culture captured different subsets of tumor clonal populations.

Duplications and deletions are not the sole form of genomic rearrangement that may induce a competitive advantage in cancer cell growth. Genomic inversions are difficult to assess through standard karyotyping and chromosome painting methods, whereas chromosomal translocations are difficult to uncover in whole-genome amplification methods, since only reads capturing the breakpoint would provide supportive evidence. To address both of these limitations, we utilized the HiC-like component of our s3-GCC libraries. Using read pairs spanning ≥50 kbp, we produced chromatin contact maps that produced clear chromatin compartmentalization signal (Figure 2i)^13^. Single cells were separated by their distal contact information via *HiCRep* and observed distinct clusters by PDCLs^29^. Notably, even at this low sequencing depth, we were able to reliably tell PDCL line sparse contact profiles apart (Figure 2j, Supplementary Figure 3). Differences between the aggregated contact maps between clusters were then used to assess unique translocation and inversion events across the sampled cells. We found that our single-cell contact data uncovers an intrachromosomal translocation between the 8.5-9.5 Mbp and 88.5-91.0 Mbp regions of chromosome 12 (Figure 2k), containing *ATP2B1*, which is commonly overexpressed in PDAC^30^ and the tumor suppressor gene *DUSP6*^31^ that is only present in PDAC-1.

Taken together, our s3 workflow represents marked improvements over the predecessor sci platform with respect to passing reads obtained per cell without sacrificing signal enrichment in the case of s3-ATAC, or coverage uniformity for s3-WGS. We also introduce another variant of combinatorial indexing workflows, s3-GCC to obtain both genome sequencing and chromatin conformation, with improved chromatin contacts obtained per cell when compared to sci-HiC. We demonstrate the utility of these approaches by assessing two patient-derived tumor cell lines with genomic instability. Our analysis reveals patterns of focal amplification for disease-relevant genes, and uncover wide-scale heterogeneity at a throughput not attainable with standard karyotyping. Additionally, we highlight the joint analysis of our protocols for uncovering the chromatin compartment disrupting effect of copy number aberrations. Furthermore, the s3 workflow has the same inherent throughput potential of standard single-cell combinatorial indexing, with the ability to readily scale into the tens and hundreds of thousands of cells by expanding the set of transposome and PCR indexes. We also expect that this platform will be compatible with other transposase-based techniques, including sci-MET^10^, or CUT&Tag^32^. Lastly, unlike sci workflows, the s3 platform does not require custom sequencing primers or custom sequencing recipes, removing one of the major hurdles that groups may face while implementing these technologies.

## Acknowledgements

We would like to thank helpful suggestions and feedback from other members of the Adey Lab as well as Jay Shendure and Cole Trapnell. This work was funded by R01DA047237 (NIH/NIDA) and R35GM124704 (NIH/NIGMS) to A.C.A; and R01MH113926 (NIH/NIMH) to B.J.O. We would also like to thank the Oregon Brain Bank for the donated biological sample used in this study.

## Author Contributions

R.M.M., D.P., F.J.S., and A.C.A. conceived the study. R.M.M. performed all s3 experiments and led all analysis under the supervision of A.C.A.; D.P. and F.Z. performed additional experiments under the supervision of F.J.S.;B.L.O. and G.G.Y. contributed to the design and analysis of chromatin conformation s3-GCC protocol and datasets; B.J.O. provided support for R.M.M. and advice on analysis; C.A.T. contributed to the analysis of cell types in the s3-ATAC datasets; J.L. generated PDCL cell lines and performed characterization of the lines under supervision of R.C.S.; J.L. and R.C.S. contributed to the analysis of PDAC s3-WGS and s3-GCC datasets. The manuscript was written by R.M.M. and A.C.A.; all authors reviewed and contributed to the manuscript.

## Methods

### PDCL propagation

Low-passage, patient-derived cell lines (PDCLs) were propagated from rapidly dissociated PDAC tumors and cultured for continuous propagation in culture medium containing ROCK inhibitor (Y-276320)^34^. Briefly, approximately 50,000 viable, disaggregated tumor cells were plated to a 35mm diameter, collagen-coated well (Gibco, A11428-02) and passaged 1:3 while subconfluent until reaching 85% confluence on a 10cm diameter dish. From a fraction of these cells, DNA was extracted to validate the presence of KRAS-G12 mutations by ddPCR (Bio-Rad, 1863506) and to validate an STR profile that matches normal leukocyte DNA from the same patient (Genetica). PDCLs exhibited morphologies consistent with epithelial tumor cells and abundant KRT expression was detected by immunocytofluorescence using the monoclonal antibodies: AE1/AE3, C-11, and Cam5.2.

### Whole Exome Sequencing and Analysis

Whole exome sequencing libraries for the patient blood sample, tumor biopsy, and PDCL were carried out by the Knight Diagnostic Research Cytogenetics Lab at OHSU. Libraries were prepared using 500 ng of fragmented gDNA using KAPA Hyper-Prep Kit (KAPA Biosystems) with Agilent SureSelect XT Target Enrichment System and Human All Exon V5 capture baits (Agilent Technologies), following manufacturer’s protocols. Sequencing was carried out using the Illumina HiSeq 2500 platform by the OHSU Massively Parallel Sequencing Shared Resource (MPSSR). Paired-end reads were aligned with *bwa mem* (v0.7.15-r1140) to GRCh38 (“hg38”,Genome Reference Consortium Human Reference 38 (GCA_000001405.2))^35^. The data was processed following the best practices workflow for the GATK pipeline (v4.1.9.0)^36^. Exome regions annotated as “protein-coding” were extracted from GenCode (v35)^37^ and used as the intervals for processing. The following commands were then used for WES data normalization and segmentation with additional options were specified: *PreprocessInvertals, CollectReadCounts, AnnotateIntervals, FilterIntervals, CreateRedCountPanelOfNormals* (using the matched blood sample as the normal, with *minimum-interval-median-percentile* set to 5.0), and finally *PlotDenoisedCopyRatios*. The output was then plotted with *ggplot2* (v3.3.2) in *R* (v4.0.0). The *geom_rect* function was used to shade the genomic region based on the relative copy number with segmentation interval, and *geom_point* was used to plot normalized bin reads.

### s3-ATAC Library Generation

Prior to sample handling, 96 uniquely indexed transposome complexes were assembled using previously-described methods^11^. Complexes were diluted to 2.5uM in a protein storage buffer composed of 50% (v/v) glycerol (Sigma G5516), 100 mM NaCl (Fisher Scientific S271-3), 50 mM Tris pH 7.5 (Life technologies AM9855), 0.1 mM EDTA (Fisher Scientific AM9260G), 1 mM DTT (VWR 97061-340), and stored at −20°C. At the time of nuclei dissociation, 50mL of nuclei isolation buffer (NIB-HEPES) was freshly prepared with final concentrations of 10 mM HEPES-KOH (Fisher Scientific, BP310-500 and Sigma Aldrich 1050121000, respectively), pH 7.2, 10 mM NaCl, 3mM MgCl2 (Fisher Scientific AC223210010), 0.1 % (v/v) IGEPAL CA-630 (Sigma Aldrich I3021), 0.1 % (v/v) Tween (Sigma-Aldrich P-7949) and diluted in PCR-grade Ultrapure distilled water (Thermo Fisher Scientific 10977015). After dilution, two tablets of Pierce(tm) Protease Inhibitor Mini Tablets, EDTA-free (Thermo Fisher A32955) were dissolved and suspended to prevent protease degradation during nuclei isolation.

For s3-ATAC tissue handling, primary samples of C57/B6 mouse whole brain were extracted and flash frozen in a liquid nitrogen bath, before being stored at −80°C. Human cortex samples from the middle frontal gyrus were sourced from the Oregon Brain Bank from a 50-year-old female of normal health status. Tissue was collected at 21 hours post-mortem and then placed in a −80°C freezer for storage. An at-bench dissection stage was set up prior to nuclei extraction. A petri dish was placed over dry ice, with fresh sterile razors pre-chilled by dry-ice embedding. 7mL capacity dounce homogenizers were filled with 2mL of NIB-HEPES buffer and held on wet ice. Dounce homogenizer pestles were held in in ice cold 70% (v/v) ethanol (Decon Laboratories Inc 2701) in 15mL tubes on ice to chill. Immediately prior to use, pestles were rinsed with chilled distilled water. For tissue dissociation, mouse and human brain samples were treated similarly. The still frozen block of tissue was placed on the clean pre-chilled petri dish and roughly minced with the razors. Razors were then used to transport roughly 1 mg the minced tissue into the chilled NIB-HEPES buffer within a dounce homogenizer. Suspended samples were given 5 minutes to equilibrate to the change in salt concentration prior to douncing. Tissues were then homogenized with 5 strokes of a loose (A) pestle, another 5 minute incubation, and 5-10 strokes of a tight (B) pestle. Samples were then filtered through a 35 µm cell strainer (Corning 352235) during transfer to a 15mL conical tube, and nuclei were held on ice until ready to proceed. Nuclei were pelleted with a 400 rcf centrifugation at 4°C in a centrifuge for 10 minutes. Supernatant was removed and pellets were resuspended in 1mL of NIB-HEPES buffer. This step was repeated for a second wash, and nuclei were once again held on ice until ready to proceed. A 10uL aliquot of suspended nuclei was diluted in 90uL NIB-HEPES (1:10 dilution) and quantified on either a Hemocytometer or with a BioRad TC-20 Automated cell counter following manufacturer’s recommended protocols. The stock nuclei suspension was then diluted to a concentration of 1400 nuclei/uL.

Tagmentation plates were prepared by the combination of 420 uL of 1400 nuclei/uL solution with 540 uL 2X TD Buffer (Nextera XT Kit, Illumina Inc. FC-131-1024). From this mixture, 8uL (∼5000 nuclei total) was pipetted into each well of a 96 well plate dependent on well schema (Figure 1b). 1uL of 2.5uM uniquely indexed transposase was then pipetted into each well. Tagmentation was performed at 55°C for 10 minutes on a 300 rcf Eppendorf ThermoMixer. Following this incubation, plate temperature was brought down with a brief incubation on ice to stop the reaction. Dependent on experimental schema pools of tagmented nuclei were combined and 2uL 5mg/mL DAPI (Thermo Fisher Scientific D1306) was added.

Nuclei were then flow sorted via a Sony SH800 to remove debris and attain an accurate count per well prior to PCR. A receptacle 96 well plate was prepared with 9uL 1X TD buffer (Nextera XT Kit, Illumina Inc. FC-131-1024,diluted with ultrapure water), and held in a sample chamber kept at 4°C. Fluorescent nuclei were then flow sorted gating by size, internal complexity and DAPI fluorescence for single nuclei following the same gating strategy as previously described^38^. Immediately following sorting completion, the plate was sealed and spun down for 5 minutes at 500 rcf and 4°C to ensure nuclei were within the buffer.

Nucleosomes and remaining transposases were then denatured with the addition 1uL of 0.1% SDS (∼0.01% f.c.) per well. 4uL of NPM (Nextera XT Kit, Illumina Inc) per well was subsequently added to perform gap-fill on tagmented genomic DNA, with an incubation at 72°C for 10 minutes. 1.5 uL of 1uM A14-LNA-ME oligo was then added to supply the template for adapter switching. The polymerase based adapter switching was then performed with the following conditions: initial denaturation at 98°C for 30 seconds, 10 cycles of 98°C for 10 seconds, 59°C for 20 seconds and 72°C for 10 seconds. The plate was then held at 10°C. After adapter switching 1% (v/v) Triton-X 100 in ultrapure H2O (Sigma 93426) was added to quench persisting SDS. At this point, some plates were stored at −20°C for several weeks while others were immediately processed.

The following was then combined per well for PCR: 16.5 ul sample, 2.5uL indexed i7 primer at 10 uM, 2.5uL indexed i5 primer at 10 uM, 3 uL of ultrapure H2O, and 25 uL of NEBNext Q5U 2X Master mix (New England Biolabs M0597S), and 0.5uL 100X SYBR Green I (Thermo Scientific S7563) for a 50 uL reaction per well. A real time PCR was performed on a BioRad CFX with the following conditions, measuring SYBR fluorescence every cycle: 98°C for 30 seconds; 16-18 cycles of 98°C for 10 seconds, 55°C for 20 seconds, 72°C for 30 seconds, fluorescent reading, 72°C for 10 seconds. After fluorescence passes an exponential growth and begins to inflect, the samples were held at 72°C for another 30 seconds then stored at 4°C.

Amplified libraries were then cleaned by pooling 25 uL per well into a 15 mL conical tube and cleaned via a Qiaquick PCR purification column following manufacturer’s protocol (Qiagen 28106). The pooled sample was eluted in 50 uL 10 mM Tris-HCl, pH 8.0. Library molecules then went through a size selection via SPRI selection beads (Mag-Bind® TotalPure NGS Omega Biotek M1378-01). 50 uL of vortexed and fully suspended room temperature SPRI beads was combined with the 50 uL library (1X clean up) and incubated at room temperature for 5 minutes. The reaction was then placed on a magnetic rack and once cleared, supernatant was removed. The remaining pellet was rinsed twice with 100 uL fresh 80% ethanol. After ethanol was pipetted out, the tube was spun down and placed back on the magnetic rack to remove any lingering ethanol. 31 uL of 10 mM Tris-HCl, pH 8.0 was then used to resuspend the beads off the magnetic rack and allowed to incubate for 5 minutes at room temperature. The tube was again placed on the magnetic rack and once cleared, the full volume of supernatant was moved to a clean tube. DNA was then quantified by Qubit dsDNA High-sensitivity assay following manufacturer’s instructions (Thermo Fisher Q32851). Libraries were then diluted to 2ng/uL and run on an Agilent Tapestation 4150 D5000 tape (Agilent 5067-5592). Library molecule concentration within the range of 100-1000bp was then used for final dilution of the library to 1 nM. Diluted libraries were then sequenced on High or Mid capacity 150 bp sequencing kits on the Nextseq 500 system following manufacturer’s recommendations (Illumina Inc. 20024907, 20024904). For greater sequencing effort, select libraries were also sequenced on a NovaSeq S2 flowcell, again following manufacturer’s recommendations (Illumina Inc. 20028315). For both machines libraries were sequenced as paired-end libraries with 10 cycle index reads and 85 cycles for read 1 and read 2.

### s3-WGS Library Generation

Prior to processing the following buffers were prepared: 50mL of NIB HEPES buffer as described above, as well as 50mL of a Tris-based NIB (NIB Tris) variant with final concentrations of 10 mM Tris HCl pH 7.4, 10 mM NaCl, 3mM MgCl2, 0.1 % (v/v) IGEPAL CA-630, 0.1 % (v/v) Tween and diluted in PCR-grade Ultrapure distilled water. After dilution, two tablets of Pierce(tm) Protease Inhibitor Mini Tablets, EDTA-free were dissolved and suspended to prevent protease degradation during nuclei isolation.

s3-WGS library preparation was performed on cell lines as follows. For patient derived PDCL cell lines, cells were plated at a density of 1×10^6^ on a T25 flask the day prior to processing. At harvest, cells were washed twice with ice cold 1X PBS (VWR 75800-986) and then trypsinized with 5mL 1X TrypLE (Thermo Fisher 12604039) for 15 minutes at 37°C. Suspended cells were then collected and pelleted at 300 rcf at 4°C for 5 minutes. For suspension-growth cell lines (GM12878), cells were pipetted from growth media and pelleted at 300 rcf at 4°C for 5 minutes.

Following the initial pellet, cells were washed with ice cold 1mL NIB HEPES twice. After the second wash, pellets were then resuspended in 300 uL NIB HEPES. Nuclei were aliquoted and quantified as described above, then aliquots of 1 million nuclei were generated based on the quantification. The aliquots were pelleted by a 300 rcf centrifugation at 4°C for 5 minutes and resuspended in 5 mL NIB HEPES. 246 uL 16% (w/v) formaldehyde (Thermo Fisher 28906) was then added to nuclear suspensions (f.c. 0.75% formaldehyde) to lightly fix nuclei. Nuclei were fixed via incubation in formaldehyde solution for 10 minutes on an orbital shaker set to 50 rpm. Suspensions were then pelleted at 500 rcf for 4 minutes at 4°C and supernatant was aspirated. Pellet was then resuspended in 1 mL of NIB Tris Buffer to quench remaining formaldehyde. Nuclei were again pelleted at 500 rcf for 4 minutes at 4°C and supernatant was aspirated. The pellet was washed once with 500uL 1X NEBuffer 2.1 (NEB B7202S) and then resuspended with 760 uL 1X NEBuffer 2.1. 40 uL 1% SDS (v/v) was added and sample was incubated on a ThermoMixer at 300 rcf set to 37°C for 20 minutes. Nucleosome depleted nuclei were then pelleted at 500 rcf at 4°C for 5 minutes and then resuspended in 50 uL NIB Tris. A 5 uL aliquot of nuclei was taken and diluted 1:10 in NIB Tris then quantified as described above. Nuclei were diluted to 500 nuclei/uL with addition of NIB Tris, based on the quantification. Dependent on experimental setup, the 420 uL of nuclei at 500 nuclei/uL were then combined with 540 uL 2X TD buffer. Following this, nuclei were tagmented, stained and flow sorted, genomic DNA was gap-filled and adapter switching was performed as described for the s3-ATAC protocol. Library amplification was performed by PCR as described above with fewer total cycles (13-15) likely due to more initial capture events per library. Libraries were then cleaned, size selected, quantified and sequenced as described previously.

### s3-GCC Library Generation

The same cultured cell line samples were harvested as described for s3-WGS library generation, and processed from the same pool of fixed, nucleosome depleted nuclei. Following quantification of nuclei, the full remaining nuclear suspensions (∼2-3 million nuclei per sample) were pooled respective of sample. Nuclei were pelleted at 500 rcf at 4°C for 5 minutes and resuspended in 90 uL 1X Cutsmart Buffer (NEB B7204S). 10 uL of 10U/uL AluI restriction enzyme (NEB R0137S) was added to each sample. Samples were then digested for 2 hours at 37°C at 300 rpm on a ThermoMixer. Following digestion, nuclear fragments then underwent proximity ligation. Nuclei were pelleted at 500 rcf at 4°C for 5 minutes and resuspended in 100uL ligation reaction buffer. Ligation buffer is a mixture with final concentrations of 1X T4 DNA Ligase Buffer + ATP (NEB M0202S), 0.01 % TritonX-100, 0.5mM DTT (Sigma D0632), 200 U of T4 DNA Ligase, diluted in ultrapure H2O. Ligation took place at 16°C for 14 hours (overnight). Following this incubation, nuclei were pelleted at 500 rcf at 4°C for 5 minutes and resuspended in 100 uL NIB HEPES buffer. An aliquot of nuclei were quantified as described previously, and were then diluted, aliquoted, tagmented, pooled, DAPI stained, flow sorted, genomic DNA was gap-filled and adapter switching was performed as described for the s3-ATAC protocol. Library amplification occurred at the same rate as the s3-WGS libraries (13-15 cycles) and libraries were subsequently pooled, cleaned, quantified and sequenced as described above.

### Computational Analysis

#### Preprocessing

The initial processing of all library types was the same. After sequencing, data was converted from bcl format to FastQ format using *bcl2fastq* (v 2.19.0, Illumina Inc.) with the following options *with-failed-reads, no-lane-splitting*, f*astq-compression-level=9, create-fastq-for-index-reads*. Data were then demultiplexed, aligned, de-duplicated using the in-house *scitools* pipeline (ref ^38^). Briefly, FastQ reads were assigned to their expected primer index sequence allowing for sequencing error (Hamming distance ≤2) and indexes were concatenated to form a “cellID”. Reads that could be assigned unambiguously to a cellID were then aligned to reference genomes. For s3-WGS and s3-GCC libraries, paired reads were aligned with *bwa mem* (v0.7.15-r1140) to hg38^35^. For s3-ATAC libraries, reads were first aligned to a concatenated hybrid genome of hg38 and GRCm38 (“mm10”, Genome Reference Consortium Mouse Build 38 (GCA_000001635.2)). Reads were then de-duplicated to remove PCR and optical duplicates by a *perl* (v5.16.3) script aware of cellID, chromosome and read start, read end and strand. From there putative single-cells were distinguished from debris and error-generated cellIDs by both unique reads and percentage of unique reads.

### s3-ATAC Analysis

#### Barnyard Analysis

With single-cell libraries distinguished, we next quantified contamination between nuclei during library generation. We calculated the read count of unique reads per cellID aligning to either human reference or mouse reference chromosomes (Figure 1C). CellIDs with ≥90% of reads aligning to a single reference genome were considered *bona fide* single-cells. Those not passing this filter (2.7%,19/687 cells for pre-tagmentation barnyard) were considered collisions. Collision rate was estimated to account for cryptic collisions (mouse cell-mouse cell or human cell-human-cell) by multiplying by two (final collision rate of 5.5%). *Bona fide* single-cell cellIDs were then split from the original FastQ files to be aligned to the proper hg38 or mm10 genomes with *bwa mem* as described above. Human and mouse assigned cellIDs were then processed in parallel for the rest of the analysis. After alignment, reads were again de-duplicated to obtain proper estimates of library complexity.

#### Tagmentation Insert Quantification

To assess tagmentation insert size, *samtools isize* (v. 1.10) was performed and plotted with *ggplot2* (v3.3.2) in *R* (v4.0.0) using the *geom_density* function (default parameters, Figure 1E). To assess library quality further, we generated tagmentation site density plots centered around transcription start sites (TSSs). We used the alignment position (chromosome and start site) for each read to generate a bed file that was then piped into the BEDOPS closest-feature command mapped the distance between all read start sites and transcription start sites (v 2.4.36)^39^. From this, we collapsed binned distances (100bp increments) into a counts table and generated percentage of read start site distances within each counts table. We plotted these data using R and *ggplot2 geom_density* function (default parameters) subset to 2000 base pairs around the start site to visualize enrichment. TSS enrichment values were calculated for each experimental condition using the method established by the ENCODE project (https://www.encodeproject.org/data-standards/terms/enrichment), whereby the aggregate distribution of reads ±1,000 bp centered on the set of TSSs is then used to generate 100 bp windows at the flanks of the distribution as the background and then through the distribution, where the maximum window centered on the TSS is used to calculate the fold enrichment over the outer flanking windows.

#### Library Complexity Analysis

To project library complexity through sequencing effort, pre-de-duplicated cellID read sets were used to build a projection as follows^8^. Reads were randomly subsampled starting at 1% of the total reads with 5% of data added in increasing increments to build a simple saturation curve per cellID. A summarized saturation curve per species was generated and plotted in *ggplot2* using the *geom_smooth* function, descripting the curves mean, median and standard error. For comparison to publicly available data sets of a matched tissue type, we focused our analysis on the mouse brain libraries. We plotted our PCR plate sequenced to 36.4% ± 17.4% unique reads/total reads for comparison to three other single-cell ATAC-seq methods which have been applied to post-natal mouse whole brain^14–16^. Data passing self-reported filters were used for comparison and plotted with *ggplot geom_boxplot* function. Welch’s two-sample T test comparisons between unique reads per cell were calculated with the *t*.*test* function in base *R* for a one-sided alternative hypothesis.

#### Dimensionality Reduction

Pseudo-bulked data (agnostic of cellID) was then used to call read pile-ups or “peaks” via macs2 (v.2.2.7.1) with option –keep-dup all^40^. Narrowpeak bed files were then merged by overlap and extended to a minimum of 500bp for a total of 292,156 peaks for human and 174,653 peaks for mouse. A scitools perl script was then used to generate a sparse matrix of **peaks × cellID** to count occurrence of reads within peak regions per cell. FRiP was calculated as the number of unique, usable reads per cell that are present within the peaks out of the total number of unique, usable reads for that cell for each peak bed file. Cells with less than 20% of reads within peaks were then filtered out. Tabix formatted files were generated using *samtools* and *tabix* (v1.7). The counts matrix and tabix files were then input into a SeuratObject for *Signac* (v1.0.0) processing^20,41^.We performed LDA-based dimensionality reduction via *cisTopic* (v0.3.0)with 27 topics for mouse cells and 24 topics for human cells^18^. The number of topics were selected after generating 25 separate models per species with topic counts of 5,10,20-30,40,50,55,60-70 and selecting the topic count using selectModel based on the second derivative of model perplexity. Cell clustering was performed with Signac *FindNeighbors* and *FindClusters* functions on the **topic weight × cellID** data frame. For *FindClusters* function call, resolution was set to 0.3 and 0.2 for human and mouse samples, respectively. The respective **topic weight × cellID** was then projected into two dimensional space via a uniform manifold approximation and projection (“UMAP”) by the function *umap* in the *uwot* package (v0.1.8, Figure 1g-h)^42^. Cis-coaccessibility networks (CCANs) were generated through the Signac wrapper of *cicero* (v1.3.4.10)^43^. Genome track plots with CCAN linkages were generated through *Signac* function *CoveragePlot* for marker genes previously described^20^. Differential accessibility between clusters in one by one, and one by rest comparisons were generated using *Signac* function *FindMarkers* using options: test.use = ’LR’, and only.pos=T, with latent.vars = ’nCount_peaks’, to account for read depth. Cell type per cluster was assigned based on genome track plots and differentially accessible sites.

#### Subclustering

After gross cell type assignment of mouse and human cell lines, human inhibitory neurons (GAD1+) clusters 3 and 4 were subset from the SeuratObject.Those 342 cells were then iteratively clustered by performing the same cisTopic, UMAP, and Signac processing with the following changes^20,42,44^. CisTopic was performed on the full set of human peaks (292,156) with those 342 subset cells. 12 Topic models were constructed (5, 10, 20-30 topics) and the 25 topic model was chosen on the second derivate of the model perplexity. A resolution of 0.5 was used in the Signac *FindClusters* on the **topic weight × cellID** call to attain 5 subclusters. Coverage plots were generated as reported above for *ADARB2* and *LHX2* . Peaks were then assigned to topics using the cisTopic *binarizecisTopics* function with argument thrP=0.975 (mean count per topic: 2429 peaks). We then performed a simple gene set enrichment analysis on human cortical inhibitory neurons and subtypes based on RNA-identified marker genes defined previously^21^.We used a Fisher’s Exact test with the function *fisher*.*test* with function alternative.hypothesis=”greater” to look for enrichment of topic-assigned peaks in marker gene bodies for inhibitory neuron subclasses relative to all topic-assigned peaks. We filtered results to those with nominal enrichment (p value ≤ 0.05) and used *ggplot geom_point* with color reflecting the reported p-value and size proportional to odds ratio to generate a bubble plot (Figure 1K).

### s3-WGS and s3-GCC Analysis

#### Quality Control

s3-WGS and s3-GCC cellIDs were initially filtered to samples with either ≥1×10^5^ or ≥1×10^6^ unique reads (PDCL and GM12878 samples, respectivley). CellIDs were split after de-duplication into single-cell bam files. They were then processed via the pipeline in the package *SCOPE* (v1.1)^23^. The genome was split into 500 kbp bins with each bin being assigned a GC content and mappability score (generated through CODEX2)^45^. Reads with a mapping quality of Q ≥ 10 were counted in bins per cellID. Bins with a mappability score < 0.9 or GC content ≤ 20% or ≥ 80% were removed (5449 bins passing filter). Additionally, cellIDs with low coverage were removed (1268 samples passing filter). Median absolute deviation (MAD) scores were calculated per cell on 500kb bins of cells passing filter as previously described^23^. Briefly, let Y_i,j_ be the raw read count for the i^th^ cellID of the j^th^ bin (from 1.. n bins). Let N_i_ be a cell-specific scaling factor (total read depth) and B_j_ be a bin-specific normalization, output as *beta*.*hat* from the function *normalize_codex2_ns_noK*. Such that

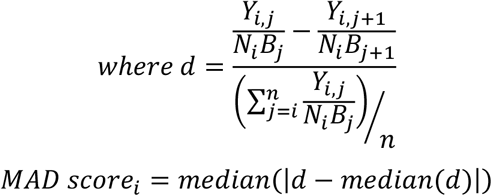

MAD scores were then plotted using the ggplot *geom_jitter* and *geom_boxplot* functions.

#### Copy Number Calling

*SCOPE* assumes diploid cells within the sample for normalization steps. To this end we used GM12878 lymphoblastoid cell line as our normal diploid samples and used an *a priori* estimate of 2.6N based on averaged PDCL karyotyping results (Figure 2c). We then used the *SCOPE* function *normalize_scope_foreach* with the following options: K=5, T=1:6 to normalize read distributions per cell. We segmented the genome into breakpoints per chromosome and inferred copy number per breakpoint per cell by *segment_CBScs* allowing for a simple nested structure of copy number changes (*max*.*ns=1*). To plot inferred copy number per cell, we used the *R* library *ComplexHeatmap* (v2.5.5) by function *Heatmap*^46^. Pairwise distance between cells was generated by Jaccard distance through the R library *philentropy* (v0.4.0)^47^ on windows categorized as “neutral” (2N), “amplified” (>2N) or “deleted” (<2N). Cells then underwent hierarchical clustering by the “ward.D2” argument in the function *hclust*. The resultant dendrogram was then cut into both 3 and 6 clades based on the two independent optimal k value searches using the *find_k* function in the R library *dendextend* (v1.14.0) given a range of 2 to 10 and 5 to 10 clusters, respectively (Figure 2i)^48^. Cells with shared clade membership were then combined into “pseudobulk” clades for higher resolution copy number calling. After combining counts data across 50 kbp bins (and filtered as described above), we had 6 clades with 154, 250, 363, 100, 268 and 133 cells, with mean reads per bin of 1207, 2442, 4662, 2071, 2700, and 9416, respectively. These pseudobulk sampled were then normalized as described above with clade 6, containing 83.45% GM12878 cells (111/133 cells) as the normal diploid sample. The genome per sample was then segmented as described above and normalized reads per bin as well as segmentation calls were plotted with *ggplot2 geom_point* and *geom_rect* functions. Select genomic locations^25^ of recurrently mutated genes were vizualized and plotted using IGV with 5 bins (250kbp) up and downstream from the transcription start sites. (Supplementary Figure 2b)^49^.

s3-GCC contact profile raw counts were generated for cellIDs passing the read count and SCOPE filters (215 cells) as follows. For initial plotting of single-cell profiles, paired-end read bam files were filtered for an insert length of ≥50kbp via *pysam*^50^ and output as upper-triangle triple-sparse format at 1mbp bin sizes. Raw contact matrices were then plotted with *R* and *ComplexHeatmap* (Figure 2j, left). Merged ensemble plots were generated by summing single-cell contact matrices generated as described above for 500 kbp bins. Following this, we performed dimensionality reduction and clustering analyses using a topic modeling approach. We treated the GCC portion of single-cell sequencing fragments (read pairs seperated by a genomic distance higher than 1kb) as traditional distal interactions. We analyzed these cells using our previously established topic model for analysis and characterization of single-cell Hi-C data^51^. In the topic modeling framework, each cell is treated as a mixture of “topics” where each topic corresponds to a set of distal interactions. The model is trained in an unsupervised manner to find the optimum number of topics that best describe the data and associates each distal interaction with a probablistic mixture of topics.

We trained a topic model using the GCC data with the default parameters in Kim et al. However, we altered one parameter, which is the range of distal interactions that are input into the model. Due to high coverage of s3-GCC assays, we opted for distal interactions that are separated by a genomic distance of 20Mb or less, as opposed to original parameter where we used interactions that are separated by distances lower than 10Mb. After training, we found that the number of topics that best describe the data is 15. We visualized cells using UMAP and found that the majority of cells from two lines cluster separately. Overall, these results validate the Hi-C like characteristics of GCC data and further show that we can capture the subtle differences in chromatin organization of the two lines.

---

**Supplementary Figure 1.**
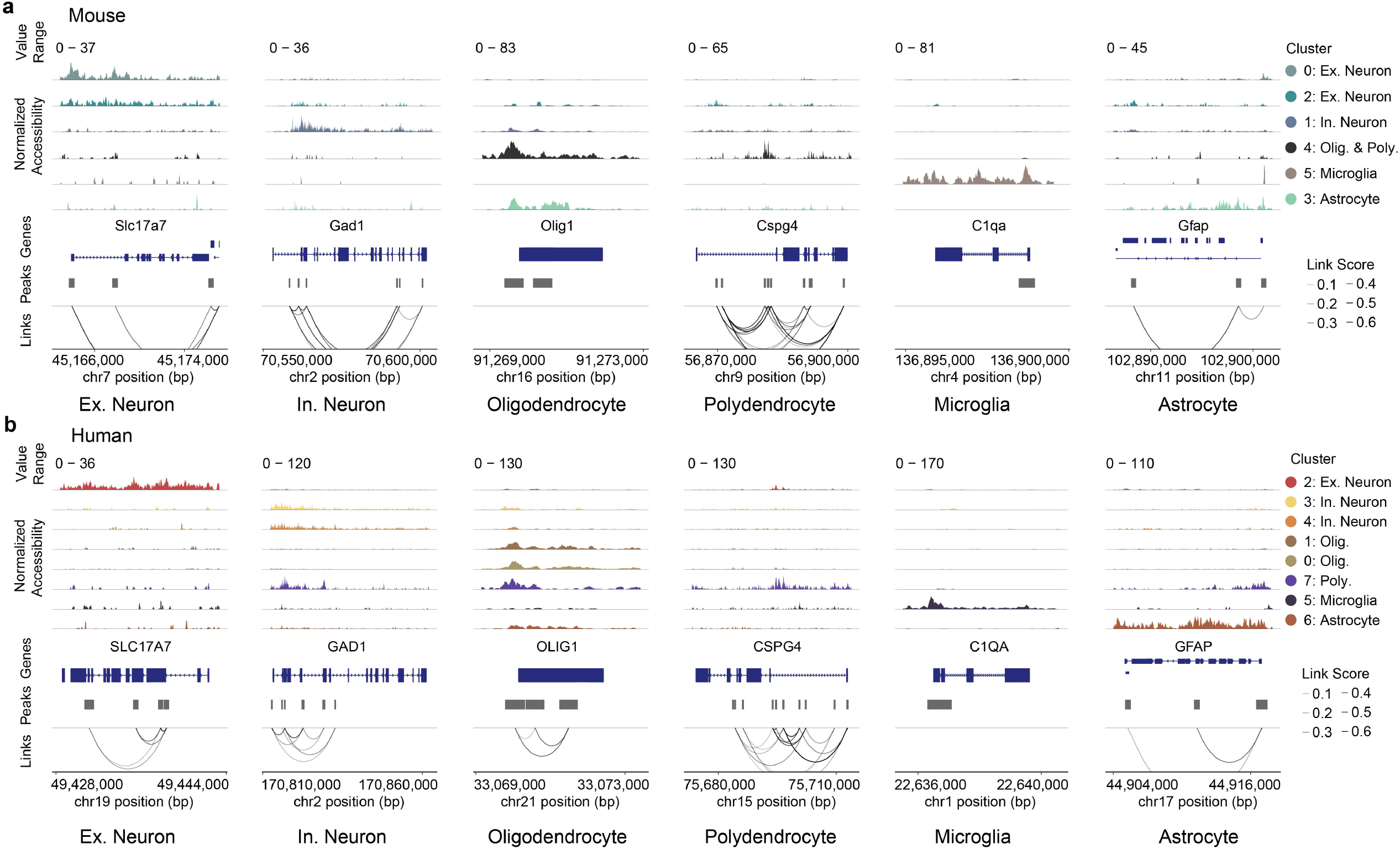
Marker sets for gross cell type discrimination in human cortex and mouse whole brain data sets. Genome coverage plots for **(a)** mouse and **(b)** human cell types. Cells are separated into tracks based on clustering.

**Supplementary Figure 2.**
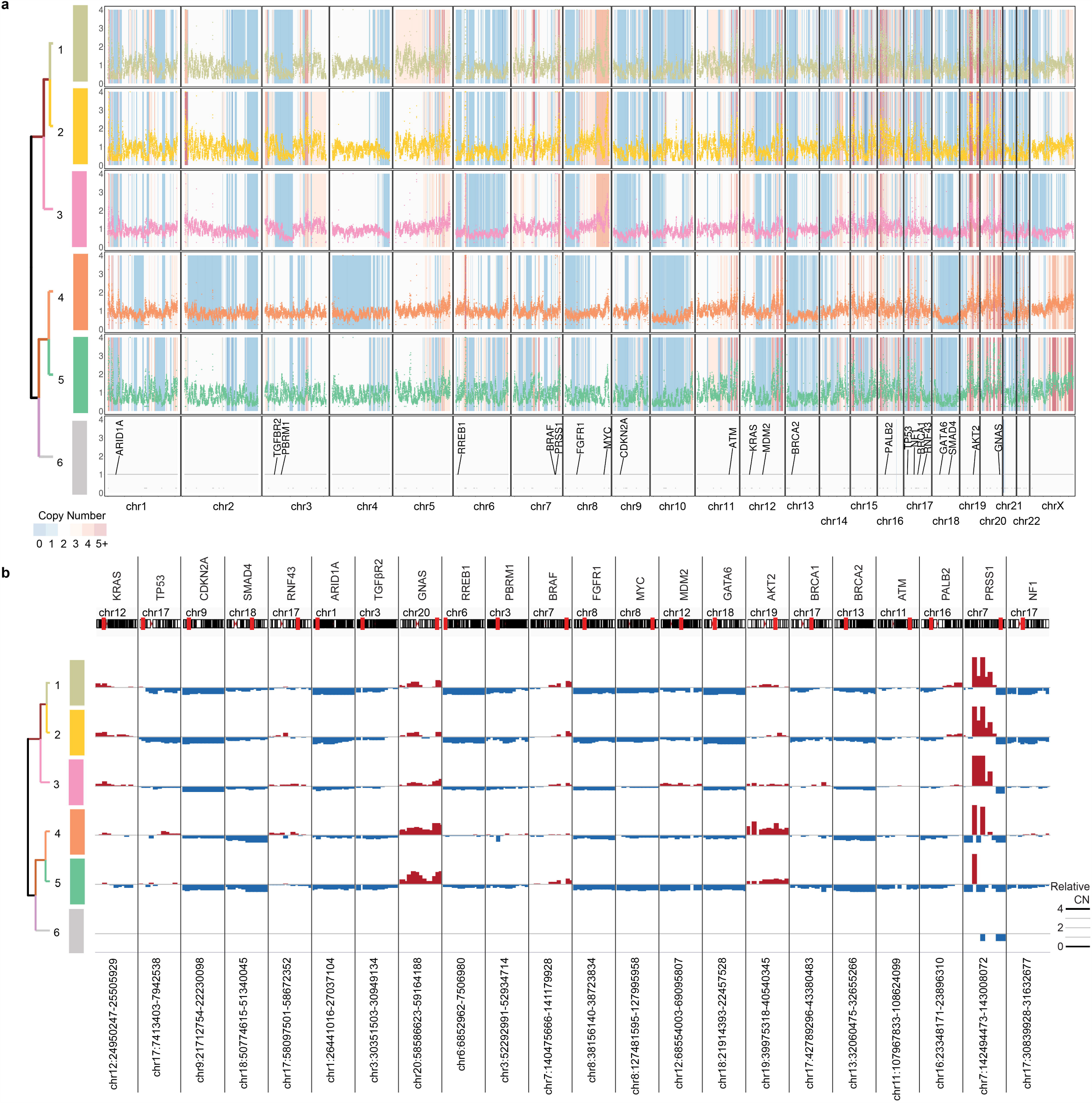
Pseudobulk whole genome copy number calling. (a) Pseudobulk genome-wide relative copy number calling. Scatter plot of reads per bin (50 kbp) separated by clade and chromosome. Clade assignment shown on left and relate to Fig 2h. Recurrently mutated gene locations in PDAC^25^ highlighted along the read count bins for clade 6. Shading of plot colored by relative copy number in genomic locus. (b) Select genes with known copy number aberrations in PDAC bar plot for relative copy number per clade. Middle line per row is copy number neutral, amplifications in red and deletions in blue.

**Supplementary Figure 3.**
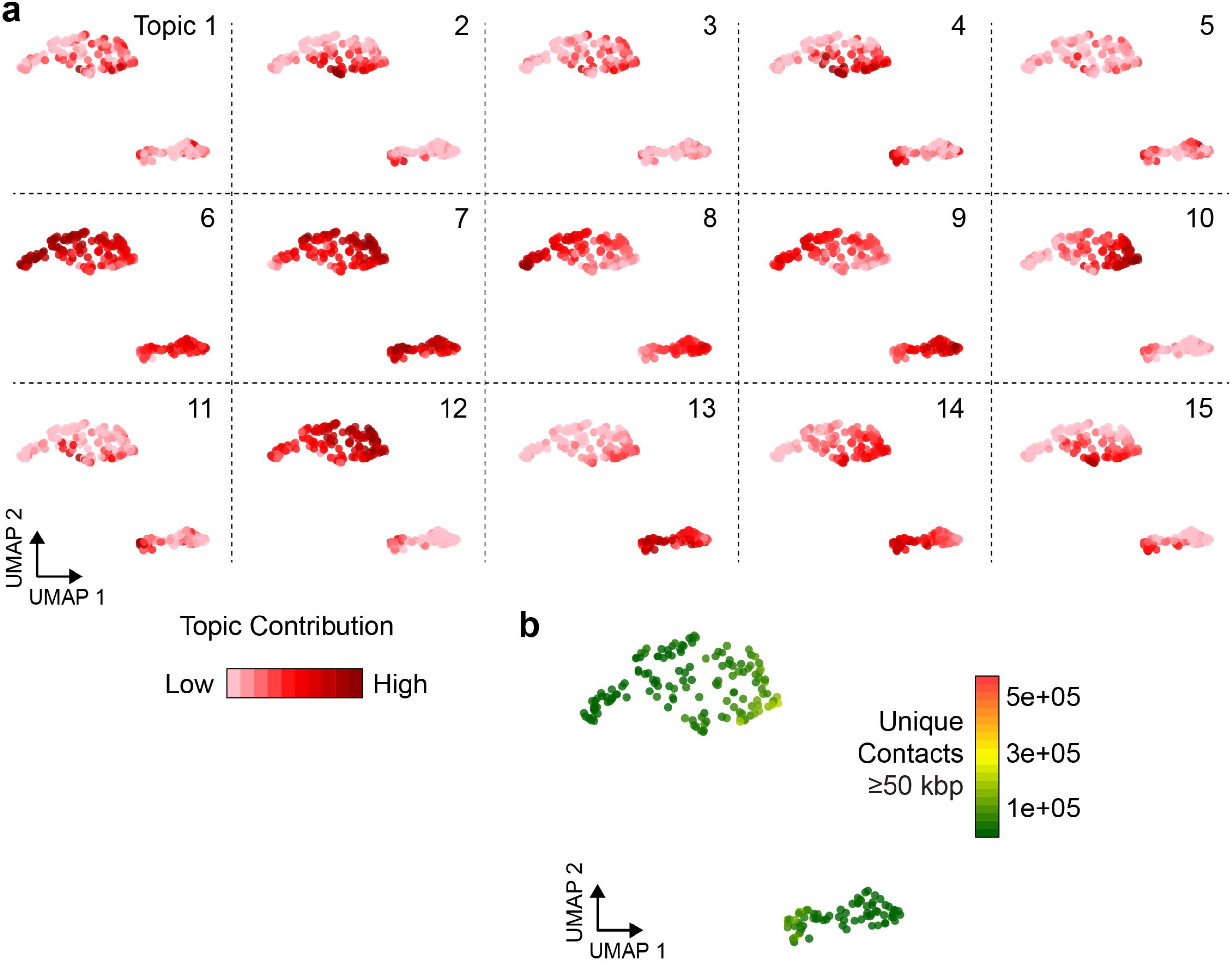
Clustering of s3GCC profiles. (a) Topic contribution scores plotted over UMAP projection of s3-GCC cells. (b) Unique contacts ≥50 kbp displayed on UMAP dimensions for quality control purposes.

